# Whole-brain computation of cognitive versus acoustic errors in music

**DOI:** 10.1101/2022.05.17.492262

**Authors:** L. Bonetti, F. Carlomagno, M. Kliuchko, B.P. Gold, S. Palva, N.T. Haumann, M. Tervaniemi, M. Huotilainen, P. Vuust, E. Brattico

## Abstract

Previous studies have evidenced how the local prediction of physical stimulus features may affect the neural processing of incoming stimuli. Less known are the effects of cognitive priors on predictive processes, and how the brain computes local versus cognitive predictions and their errors. Here, we determined the differential brain mechanisms underlying prediction errors related to high-level, cognitive priors for melody (rhythm, contour) versus low-level, local acoustic priors (tuning, timbre). We measured with magnetoencephalography the mismatch negativity (MMN) prediction error signal in 104 adults having varying levels of musical expertise. We discovered that the brain regions involved in predictive processes for local priors were primary and secondary auditory cortex and insula, whereas cognitive brain regions such as cingulate and orbitofrontal cortices were recruited for melodic errors in cognitive priors. The involvement of higher-level brain regions for computing cognitive errors was enhanced in musicians, especially in cingulate cortex, inferior frontal gyri, and supplementary motor area. Overall, the findings expand knowledge on whole-brain mechanisms of predictive processing and the related MMN generators, previously mainly confined to the auditory cortex, to a frontal network that strictly depends on the type of priors that are to be computed by the brain.

## Introduction

According to predictive coding theory, audition is an active process where models of expectations for the incoming sounds are constantly updated based on expectations (also termed priors) when errors occur ^1,2^. Recent neuroimaging studies recorded the mismatch negativity (MMN) with electroencephalography (EEG) or magnetoencephalography (MEG) ^3,4^ providing empirical support for the theory in relation to auditory processing, since MMN indexes predictive coding errors of acoustic features and demonstrate the existence of ascending, forward connections in the auditory cortex conveying prediction errors, which correspond to the ‘new’ information conveyed by the external stimuli that cannot be predicted. MMN studies also demonstrated the presence of backward connections from higher-order areas of the auditory cortex to predict activity in lower-order areas ^5,6,7,8^.

However, most studies measured the MMN for simple acoustic feature errors, analyzing the MEEG sensor signal and parameters. Only a minority of studies have provided a clear reconstruction of the neural sources. These studies returned a network of active brain areas that were mainly localized in the auditory cortex and especially in Heschl’s gyrus, superior, and middle temporal gyri ^9–12^. Additional, weaker generators of the MMN were localized in the inferior frontal cortex and cingulate gyrus ^9–11^. Functional magnetic resonance imaging (fMRI) studies confirmed the involvement of superior temporal gyrus and right inferior and middle frontal gyri in the generation of the MMN ^13,14^. Taken together, the current literature supports the hypothesis that auditory cortex is the main generator of the MMN elicited in response to errors of acoustic priors, with frontal generators possibly responsible for the process of involuntary “attention switching” and prior updating ^15–19^.

Within this framework, music listening is a peculiar case: while processing music the brain constantly predicts lower-level acoustic features using knowledge (priors) accumulated from life-long exposure to all kinds of sounds while, at the same time, integrating them with music-specific priors from exposure to a specific musical culture, allowing us to spot those changes and mistakes (e.g., in tonality, harmony, transposition, rhythm) that make music either interesting and pleasurable or, conversely, boring and dissonant ^6,20,21^. How this process, indexed by the MMN prediction error signal, is enacted in the different brain areas, beyond the primary auditory cortex, is, thus far, an open research question.

Most of MMN studies of musical sound processing, including the fMRI ones, utilized simple auditory oddball paradigms, where the acoustic features inserted in sequences of coherent sounds (e.g., pitch, rhythm, location, timbre) are broken by sudden, infrequent deviant sounds ^22,23^. These oddball paradigms eliciting the MMN response have allowed us to study automatic predictive processes for sounds that rely on feedforward and backward projections from and to the auditory cortex, not requiring the intervention of additional attentional resources ^4^. However, these oddball sequences have little resemblance with the variety of sounds and sound features that are encountered in music, limiting our understanding of whether predictions can be made at that sensory level also for the more cognitive features of sounds that are essential in music ^24^.

To this end, the newer “multi-feature” paradigm ^25,26,27^ introduces a deviation in a single feature into every second sound of a musical pattern, allowing for the recording of several MMNs of prediction errors. In a musical version of this paradigm, six deviants were used (pitch, slide, duration, timbre, location or intensity), obtaining reliable MMNs ^28–32^. Similarly, in the latest “MusMelo” paradigm, six deviants are inserted in a loop of one elaborated musical melody ^33,34^, crucially including two distinct categories of deviants: cognitive or high-level deviants and acoustic or low-level ones. Cognitive deviants refer to changes in the melodic line (melodic contour) of the melody, altering the meaning of the music since they give rise to a varied version of the original melody. Conversely, acoustic deviants sound merely like “mistakes” during the musical performance without producing any actual change of the melodic line. Hence, the MusMelo paradigm offers a unique possibility of measuring the neural indexes of cognitive versus acoustic priors and their related prediction error signals, and of locating the subservient neural sources.

To summarize, much is known on the MMN prediction error signal and its neural substrate in the auditory cortex. However, there is not definitive consensus on the role of frontal MMN generators, especially in the music perception domain. In this study, we wished to determine the predictive processes in the whole brain that are responsible for generating error signals in music, when the error is computed against acoustic versus cognitive priors. To this goal, we investigated in a large sample of over 100 participants the neural sources of acoustic versus cognitive errors of musical melodies. We hypothesized that we would observe stronger frontal generators for cognitive deviants, and increased responses in the auditory cortex to acoustic deviants.

Finally, MMN has been repeatedly connected to cognitive abilities ^35,36^ and musicianship, musical learning and cognitive abilities ^29,37–40^. For instance, Putkinen and colleagues ^34^ showed an enhanced MMN in children exposed to musical training, especially for melody modulation, mistuning and timbre. Interestingly, such differences were not existent before exposure to the musical training. Investigating the relationship between MMN and musical training, Kliuchko and colleagues ^41^ discovered an overall stronger MMN to timbre, pitch and slide for jazz compared to classical musicians and non-musicians and amateurs. Similarly, Tervaniemi and colleagues ^42^ showed that MMNs were enhanced for tuning deviants in classical musicians, for timing deviants in classical and jazz musicians, and for transposition deviant in jazz musicians. Moreover, fMRI evidence has consistently demonstrated how musical expertise refines cognitive priors making them accurate and sophisticated, for instance, allowing musicians to notice subtle harmonic changes and their action observation areas and higher-order frontal areas to be activated during mere music listening ^43,44^. For all these reasons, in this study we also assessed whether the MMN generators to cognitive and acoustic deviants were modulated by the participants’ musical expertise.

## Methods

### Participants

Participants were volunteers recruited with fliers in social media or posted in academies and universities, and they were compensated for the time spent in the lab with vouchers that could be used for culture and sport activities (e.g. museums, concerts, swimming pools, etc.). Prior to the beginning of the experiment, participants filled in the informed consent. The experimental procedures, included in the wide research protocol named “Tunteet” (“Emotions” in Finnish), complied with the Declaration of Helsinki – Ethical Principles for Medical Research, and were approved by the Ethics Committee of the Hospital District of Helsinki and Uusimaa (approval number: 315/13/03/00/11, dated 11^th^ March 2012). The “Tunteet” protocol included 1 to 3 experimental sessions (depending on participants’ availability). Besides the paradigm included for this study, other paradigms were presented to the participants, however, never exceeding 60 minutes of MEEG recordings and three hours of time spent at the Biomag laboratory (including welcoming, preparation, instructions, questionnaires and forms filling, and dismissal). MRI recordings were conducted in another day separated by maximum two weeks from MEG recordings. Other MEG and behavioral findings with the same participants are reported in Kliuchko and colleagues ^41^, Haumann and colleagues ^45^, Criscuolo and colleagues ^35^, and Bonetti and colleagues ^30,31^. The current dataset based on the Musmelo paradigm has, however, never been reported in a paper.

The study comprised 104 volunteers: 44 males and 60 females (age range: 18 – 51 years old, mean age: 28.24 ± 7.92 years). All participants declared to be healthy and reported no current or previous drug nor alcohol abuse. In addition, they were not under any kind of medication, they did not have any neurological or psychiatric disorder, and declared to have normal hearing. Finally, their educational, economic, and social statuses were homogeneous, as studied and reported in Criscuolo and colleagues ^35^.

Since musicianship has been connected to modulation of MMN responses ^17,39,50–52^, we recruited participants with different levels of musical expertise. Specifically, the average formal musical training received by our participants was 5.88 ± 7.12 years (ranging from 0 to 28 years of musical training). Indeed, our samples comprised musicians who obtained a professional musical education or graduated from Sibelius Academy and University of Helsinki, amateur musicians who had only few years of formal musical training, and non-musicians.

### Experimental design and stimuli

To detect the brain predictive responses to cognitive and acoustic deviants, we used the Melodic Multifeature paradigm (MusMelo) introduced by Tervaniemi and colleagues ^33^ and Putkinen and colleagues ^34^ while participants’ brain activity was recorded by means of magnetoencephalography (MEG).

The MusMelo paradigm consisted of brief recursive melodies composed by the author Minna Huotilainen. These melodies were played with the standard timbre correspondent to digital piano tones (McGill University Master Samples) and followed typical Western tonal musical harmonies and configurations.

The melodies started with a triad (duration of 300 ms), followed by four tones of different length, plus an ending tone. To be noted, a 50-ms gap was always present between successive tones, while the ending tone was of 575 ms duration. Additionally, a 125 ms gap between each melody was inserted. Thus, one melody lasted for 2100 ms in total. Such melodies were presented for 15 min in a looped, recursive manner.

Within these repeated melodies, six different deviants (changes) were inserted. Importantly, they were divided into low-level, acoustic deviants and high-level, cognitive deviants.

The key difference between the two categories of deviants is that low-level, acoustic deviants did not alter the melodic contour of the musical stimuli, but introduced acoustic mistakes (e.g., small variations in pitch or rhythm that were perceived as mistakes and not drastic changes of the melodies). Conversely, high-level, cognitive deviants operated a profound change in the melodies that were perceived as proper variations. One melody could contain several changes, as illustrated in **Figure 1**.

**Fig. 1.**
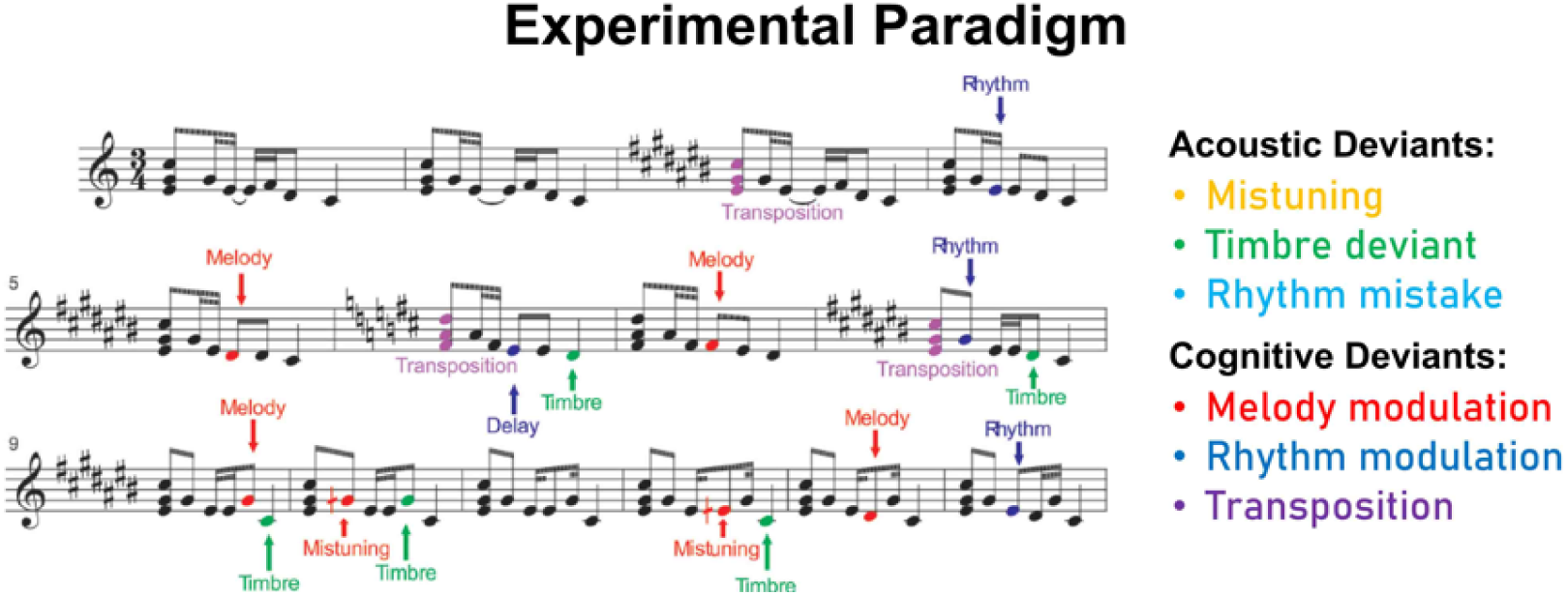
Melodic multi-feature (MusMelo) paradigm. Graphical depiction of the MusMelo paradigm, consisting of brief recursive melodies played consecutively in a loop. In these melodies, six different deviants have been inserted. The deviants belonged to two categories: acoustic deviants (mistuning, timbre, rhythm mistake) and cognitive deviants (melody modulation, rhythm modulation, transposition).

As follows, we provided details on the acoustic deviants:

1. *Mistuning* (half of a semitone upwards, up to 3% of the fundamental frequency of the sound). It occurred in the 14% of the melodies and could happen in the first, second or fourth tone of the melody.
2. *Timbre deviant* (flute timbre instead of the standard piano timbre). It occurred in the 8% of the melodies and could happen in the first, third or fourth tone of the melodies.
3. *Timing delay* (100 ms silent gap). It occurred in the 8% of the melodies. It could happen in the first, second or third tone.

Conversely, these were the characteristics of the cognitive deviants:

1. *Melody modulation* occurred in the 12% of the melodies. It consisted of a pitch change of the third or fourth tone. It endured until a new melody modulation was introduced.
2. *Rhythm modulation* occurred in the 7% of the melodies and could happen in the second or third tone. There were two possible alternatives for rhythm modulation, either a short tone was replaced by a long tone (tone lengthening) or a long tone was replaced by a short one (tone shortening).
3. *Transposition* occurred in the 16% of the melodies and could occur in the first triad. In this case, after introducing the chord transposition the following melodies kept the converted key until a new chord transposition was presented.

Thus, all cognitive deviants became the repeated form of the melody in its subsequent presentations. This system has been previously called roving-standard fashion ^46^. Finally, all cognitive deviants were musically plausible, both when the change involved the melodic contour and the rhythm contour.

The stimuli were presented using Presentation software (Neurobehavioural Systems, Berkeley, CA). In a separate session, the structural images of participants’ brain were acquired by using magnetic resonance imaging (MRI).

### Data acquisition

MEG data was collected at the Biomag Laboratory of the Helsinki University Central Hospital. The measurements were conducted in a magnetically shielded room (ETS-Lindgren Euroshield, Eura, Finland) with Vectorview™ 306-channel MEG scanner (Elekta Neuromag®, Elekta Oy, Helsinki, Finland). The MEG scanner had 102 sensor elements. Specifically, it had 102 orthogonal pairs of planar gradiometer SQUID sensors and 102 axial magnetometer SQUID sensors. We placed electrodes above and below the left eye and close to the external eye corners on both sides of the face of the participants to record horizontal and vertical eye movements. Furthermore, we recorded the continuous head position of the participants by using the head position indicator (HPI) coils that were placed on the forehead and behind the ears of participants. Moreover, for each participant we recorded the fiducial points corresponding to nasion and to the prearicular anatomical landmarks by using the Isotrack 3D digitizer (Polhemus, Colchester, VT, USA). The HPI coils and fiducial points were necessary to perform co-registration between MEG and MRI data at a later stage of analysis. Finally, the MEG data was registered with a sampling rate of 600 Hz.

The recorded MRI data was the structural T1, required for the source reconstruction of the MEG signal. The MRI scanning was conducted using a 3T MAGNETOM Skyra whole-body scanner (Siemens Healthcare, Erlangen, Germany), plus a standard 20-channel head-neck coil. The measurements were done at the Advanced Magnetic Imaging (AMI) Centre (Aalto University, Espoo, Finland). Details of the T1-weighted structural images are reported as follows: 176 slices; matrix = 256×256; field of view = 256×256 mm; pulse sequence = MPRAGE; slice thickness = 1 mm; interslice skip = 0 mm. Later in the analysis pipeline, we co-registered each individual T1-weighted MRI scan to the standard MNI brain template through an affine transformation. Then, we referenced such image to the MEG sensors space by employing the Polhemus head shape data and the three fiducial points collected prior to start the MEG recording.

### Data pre-processing

We preprocessed the raw MEG sensor data by using the signal space separation solution implemented in MaxFilter ^47^ which attenuated the interference originated outside the scalp. Afterwards, we converted the data into the SPM format and further analysed it in Matlab (MathWorks, Natick, Massachusetts, United States of America) by employing OSL (OHBA Software Library), a freely available toolbox that relies on a combination of FSL ^48^, Fieldtrip ^49^, SPM ^50^, as well as in-house-built functions.

First, a few segments of the data contaminated by large artifacts were removed after visual inspection. Second, we corrected the brain data for the interference of eyeblinks and heart-beat artefacts by using independent component analysis (ICA). This procedure decomposed the original signal in independent components. Then, we identified and discarded the components that picked up the eyeblink and heart-beat activities. Finally, we rebuilt the signal by using the remaining components ^51^. After the preprocessing steps, we epoched the signal in 4130 trials (one for each sound) lasting 700 ms each (with 100ms of pre-stimulus time that was used for baseline correction). To be noted, in a few cases the number of trials was lower than 4130. This happened when a few segments of the data were previously discarded due to the presence of large artefacts.

### MEG sensor analysis

Although our focus was on the MEG source reconstructed brain data, a first analysis on MEG sensors data was computed, in accordance with state-of-the-art guidelines about best practice in MEG analysis ^52^.

Thus, according to a large number of MEG and electroencephalography (EEG) task-related studies ^52–54^, we averaged the trials over conditions, and we combined planar gradiometers by sum-root square. Then, we assessed whether the deviant stimuli elicited a clear MMN signal by contrasting the brain responses to our six categories of deviants against the standard stimuli. Since this contrast has been done for each deviant (six), each time point (182, ranging from 0 to 300 ms from the onset of the deviant stimuli), and each MEG combined gradiometer channel (102), we have corrected for multiple comparisons by using Bonferroni correction and thus lowering the *p-value* to 4.5e-07 (.05 / (6 * 182 * 102)). In this analysis, we used combined gradiometers only because of their better signal-to-noise ratio than magnetometers when performing analysis on the MEG sensor level ^52^. The results showed that the MMN was strongly elicited among all deviants, are illustrated in **Figure 2**. The detailed statistical results showing significant time-points and channels for each deviant are reported in **Table ST1**.

**Fig. 2.**
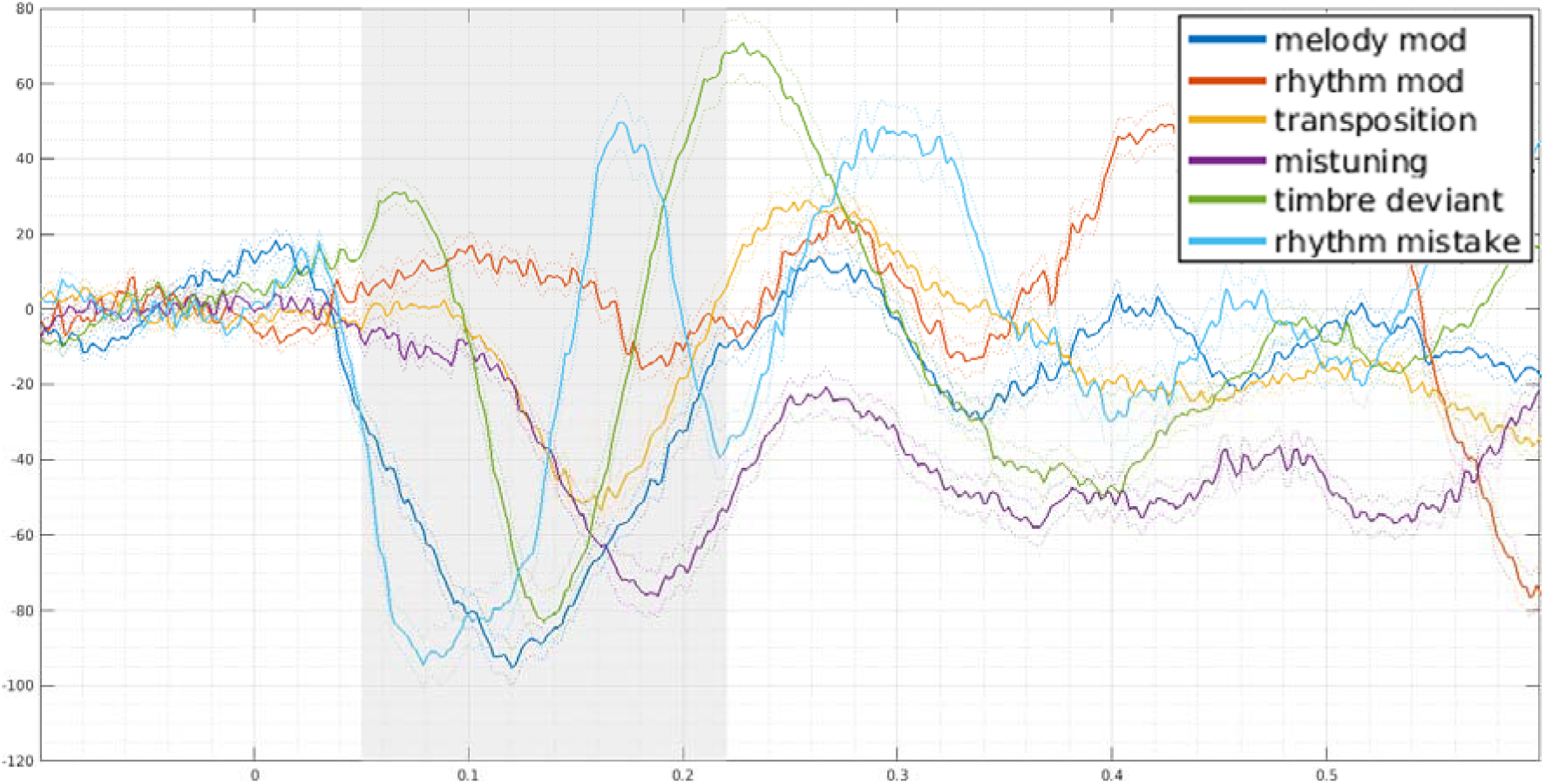
MMN to all deviants (MEG channel 1341). Waveform depicting the MMN responses (deviant minus standard) to the six deviants occurring in the MusMelo paradigm (melody modulation, rhythm modulation, transposition, mistuning, timbre, and rhythm mistake). The time series were recorded at the MEG magnetometer channel 1341, which is a typical channel shown in MMN studies. Dash lines show the standard error. The grey area highlights the different peaks of the MMN to the six deviants included in the study. X-axis shows time (in seconds), while y-axis amplitude of the signal in fT.

### Source reconstruction

We reconstructed the neural sources of the brain activity recorded on the scalp by the MEG channels, applying the widely adopted procedure named beamforming. Here, we used the OSL implementation consisting of a local-sphere forward model and a beamformer approach as the inverse method ^55–57^. The local-sphere forward model considers the MNI-co-registered anatomy as a simplified geometric model, and it fits a sphere separately for each sensor ^58^. Then, the beamforming employs a different set of weights sequentially applied to the source locations to isolate the contribution of each source to the activity recorded by the MEG channels for each different time point. In our procedure, we used a three-dimensional eight-mm grid which resulted in a brain parcellation of 3559 dipoles (sources) and both magnetometers and (non-combined) planar gradiometers.

### Neural sources of MMN peaks

We computed an independent GLM sequentially for each time point at each dipole location, where we contrasted each deviant category against the standard stimuli. This procedure, computed independently for each participant, allowed us to detect the contrast of parameter estimates (COPEs) for the brain activity specifically associated with the detection of the deviant stimuli (i.e. the MMN in source space). These results were then submitted to a second-level (group) analysis, using one-sample t-tests with spatially smoothed variance obtained with a Gaussian kernel (full-width at half-maximum: 50 mm).

Although the analysis was computed for each time-point in the epoch, we were only interested in the brain sources of the MMNs peak. Thus, after detecting the peak MMN activity independently for each deviant, we extracted and averaged the group-level results around the MMN peak (considering a small time-window of ± 25ms around the MMN peak). This procedure returned the strength of the MMNs to the six deviants for each brain dipole. To correct for multiple comparisons, we performed a cluster-based permutation test with 5000 permutations which allowed us to isolate the clusters of brain activity underlying the generation of the MMNs. Since we computed six tests (one for each deviant), we have used an α level of .0017 (.05/6), corresponding to a cluster forming threshold *t*-value = 3.3.

### MMNs to cognitive versus acoustic deviants

After detecting the sources of the brain signals underlying the MMNs peak, we performed a further analysis to assess whether such sources differed according to the category of deviants. Specifically, we were interested in assessing whether cognitive deviants (transposition, melody modulation and rhythm modulation) elicited an MMN with different brain sources than acoustic deviants (mistuning, timbre, rhythm mistake). Thus, first we averaged together the neural activity of the three deviants forming the two categories. We conducted this procedure independently for each participant. Second, we computed a t-test for each brain dipole comparing the brain activity underlying cognitive versus acoustic deviants. Finally, to correct for multiple comparisons, we performed cluster-based Monte-Carlo simulations (MCS) ^53,59–61^. Specifically, the MCS consisted of detecting the spatial clusters of significant dipoles (dipoles whose test had a *p-value* lower than the MCS α level) in the original data and assessing whether they were significant or occurred by chance. First, spatial clusters were identified in the original data. Then, we permuted the original data and detected the clusters in this new permuted set of brain values. We computed this procedure 1000 times, obtaining a reference distribution of cluster sizes detected for each permutation. Finally, the original cluster sizes were compared to the reference distribution and considered significant if they were bigger than 99.9% (MCS *p-value* of .001) of the permuted cluster sizes. In this case, we computed two MCS, one for the significant dipoles where cognitive deviants were stronger than acoustic ones, and another one for the dipoles where the acoustic deviants were stronger than the cognitive ones. Remarkably, while cognitive versus acoustic deviants returned a significant cluster only with a standard cluster-forming threshold *p-value* = .05, acoustic versus cognitive deviants returned a significant cluster even when lowering the cluster-forming threshold *p-value* to 1.0e-04, indicating a very large significant difference. Details of the outcomes of these analyses are reported in the Results section.

### Cognitive, acoustic deviants and musicianship

The last step of our analysis pipeline was to assess whether there was a relationship between musical expertise and the brain areas activated during the perception of cognitive and acoustic deviants. Thus, we computed Pearson’s correlations for each brain dipole between the participants’ years of music playing and their brain activity underlying deviant detection. Afterwards, we corrected for multiple comparisons employing an MCS analogous to the one described above. In this case, since we computed two independent MCS analyses, one for the cognitive and one for the acoustic deviants, we used a cluster-forming threshold *p-value* = .01 and an MCS *p-value* = .001.

## Results

### Experimental design and MMNs detection

Our study had three main aims: reconstructing the neural sources of the deviants inserted in a melodic multifeature paradigm (i), assessing whether such neural sources differed across cognitive and acoustic deviants (ii), investigating the relationship between the neural sources of cognitive and acoustic deviants and the musical expertise of the participants.

We employed the Melodic Multifeature paradigm (MusMelo), which was introduced by Tervaniemi and colleagues ^33^ and Putkinen and colleagues ^34^ and consists of a series of deviants breaking cognitive (transposition, melody modulation and rhythm modulation) or acoustic (mistuning, timbre, and rhythm mistake) musical features. Thus, it is the ideal paradigm to assess whether the MMNs neural sources vary depending on the characteristic of the deviants (cognitive versus acoustic). To address our research questions, we presented our 104 participants with the MusMelo paradigm while we collected their brain activity using MEG.

Although our focus was on the neural sources of the MMNs elicited by the six deviants of the MusMelo, prior to computing the analysis in MEG source space, we verified that we had obtained a reliable MMN signal on the MEG sensors. We computed one-sample t-tests for each MEG-combined gradiometer channel (102), each time point (182, ranging from 0 to 300 ms after the onset of the stimuli), and each deviant, comparing the brain response to the deviant versus the standard stimuli. We corrected for multiple comparisons by using Bonferroni correction which resulted in an adjusted *p-value* of 4.5e-07 (.05 / (6 * 182 * 102)). Our results showed that the MMNs were clearly identified among several MEG channels and time points (*p* < 4.5e-07), as illustrated in **Figure 2** and reported in detail in **Table ST1**.

### Neural sources of MMN peaks

After verifying the reliability of our paradigm in detecting clear MMN signals, we reconstructed the sources of the neural signal by combining the MEG and MRI data of each participant. Specifically, as widely done in the field, we used a local-sphere forward model and a beamformer approach as the inverse method (see Methods for details). Our procedure returned a time series describing the strength of the neural signal over time for each category of stimuli (deviant and standard) and for each of the reconstructed 3559 brain sources (dipoles). Then, we computed a first-level analysis (independently for each participant) by computing a GLM sequentially for each time point at each dipole location, where we contrasted each deviant category against the standard stimuli. These results were submitted to a second-level (group) analysis, using one-sample t-tests with spatially smoothed variance obtained with a Gaussian kernel (full-width at half-maximum: 50 mm). Although the analysis was computed for each time point in the epoch, we were only interested in the brain sources of the MMNs peak. Thus, first we detected the peak MMN activity independently for each deviant. Second, we extracted and averaged the group-level results around the MMNs peak (considering a time window of ± 25 ms around the MMN peak). Third, we corrected for multiple comparisons using a cluster-based permutation test ^65^ with 5000 permutations which allowed us to isolate the significant clusters of brain activity underlying the generation of the MMNs. Since we computed six tests (one for each deviant), we have used an α level of .0017 (.05/6), corresponding to a cluster forming threshold *t*-value = 3.3.

As depicted in **Figure 3** and reported in detail in **Table ST2**, these analyses (*p* < .0017) returned a main involvement of the primary and secondary auditory cortices, especially for timbre, rhythm mistake, melody modulation, and mistuning. Remarkably, medial cingulate gyrus and hippocampal regions were also strongly activated by the presentation of the deviant stimuli. This result was particularly evident for melody and rhythm modulation, rhythm mistake, and timbre. Finally, transposition, which is a rather cognitive and complex deviant, elicited activity mainly localized in the anterior part of the cingulate and in the inferior frontal gyrus.

**Fig. 3.**
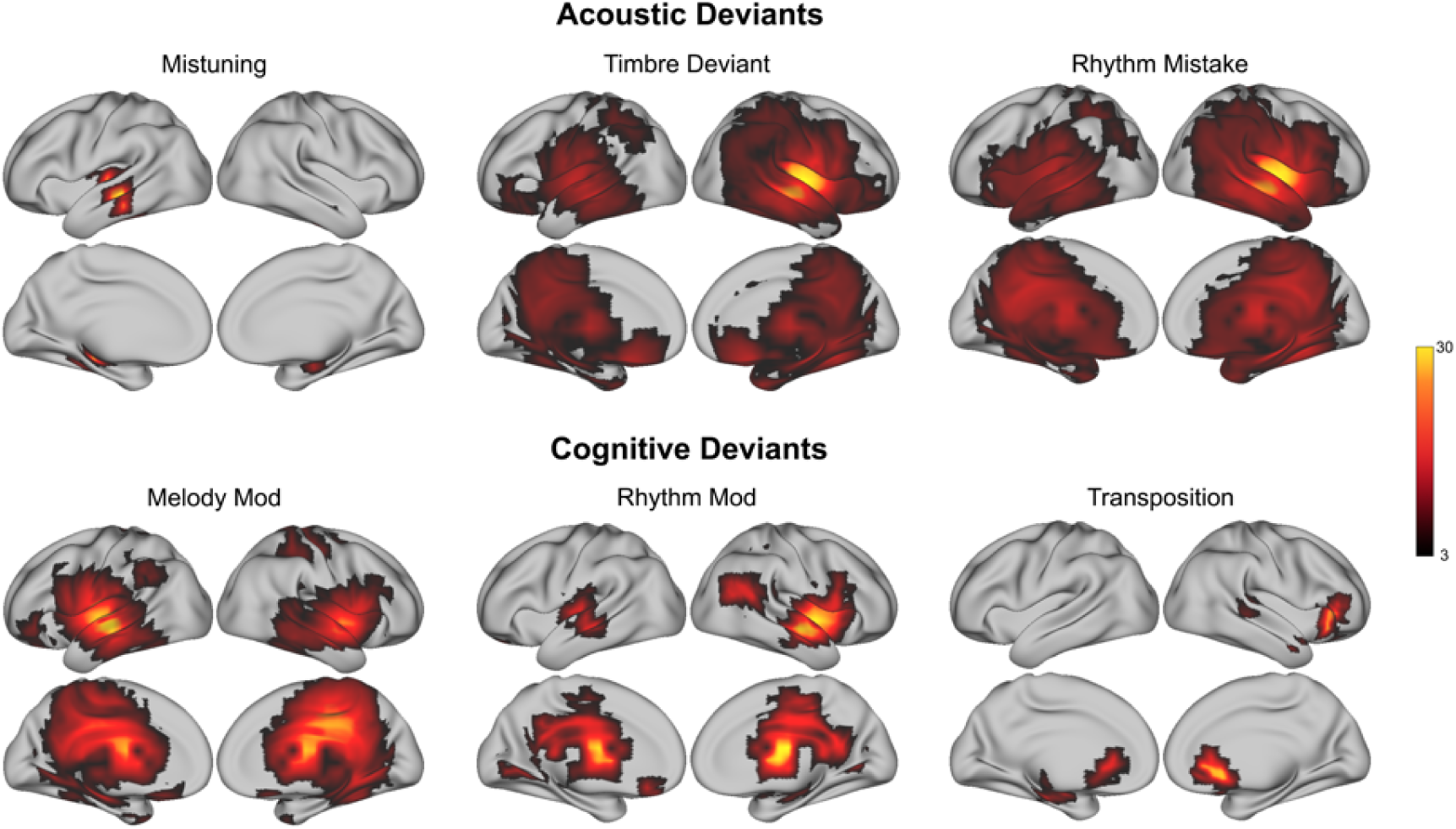
Brain sources of the MMN to all deviants. Brain sources of the MMN to all deviants depicted in brain templates. The colorbar indicates t-values obtained by contrasting the brain response to deviant versus standard sounds. The top row illustrates acoustic deviants, while the bottom row depicts cognitive deviants. Overall, acoustic deviants show strong activity in the auditory cortex, while cognitive deviants highlight the contribution of cingulate and frontal brain areas to the generation of the MMN.

### MMNs to cognitive versus acoustic deviants

After detecting the sources of the brain signals underlying the MMNs peak, we performed a further analysis to assess whether these sources differed when comparing cognitive (transposition, melody modulation, and rhythm modulation) versus acoustic (mistuning, timbre, and rhythm mistake) deviants. Thus, first we averaged together the neural activity of the three deviants in each category. Second, we computed a t-test for each brain dipole comparing the brain activity underlying cognitive versus acoustic deviants. Finally, to correct for multiple comparisons, we performed cluster-based MCS (MCS *p-value* < .001).

When using a cluster-forming threshold *p-value* < .05 (see Methods for details), we identified a small, but significant cluster of activity where cognitive deviants had a stronger neural signal than acoustic ones. This cluster was localized in the medial cingulate gyrus.

Remarkably, when computing the MCS to identify the clusters where the brain activity was stronger for acoustic versus cognitive deviants, we observed a large cluster which was largely significant (cluster forming threshold *p-value* < 1.0e-04). This cluster mainly originated in the right primary auditory cortex, but extended to secondary auditory cortex, insula, frontal operculum, and hippocampal regions (**Figure 4A** and **Table ST3**).

**Fig. 4.**
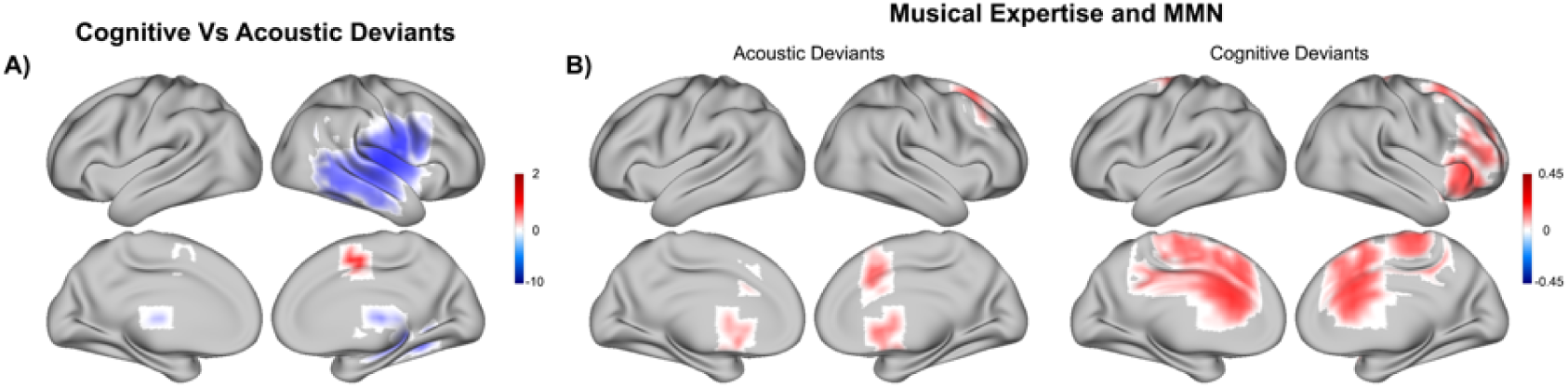
MMN to acoustic versus cognitive deviants and musical expertise. **(A)** Depiction in brain templates of the contrast between cognitive versus acoustic deviants. The colorbar shows the t-values emerged from the contrast. **(B)** Depiction in brain templates of the correlation between musical expertise and MMN to acoustic (left) and cognitive (right) deviants. The colorbar shows the r-value obtained from the correlations.

### Cognitive, acoustic deviants and musicianship

Finally, we wished to assess whether there was a relationship between musical expertise and the neural sources of the MMNs elicited by cognitive and acoustic deviants. Thus, we computed Pearson’s correlations between the participants’ years of music playing and their brain activity underlying deviant detection. This analysis was computed for each brain source originating the peak of the MMNs. We corrected for multiple comparisons employing an MCS analogous to the one described above (cluster-forming threshold *p-value* < .01 and MCS *p-value* < .001).

This analysis showed significant clusters of positive correlations between musical expertise and neural response to deviants (**Figure 4B** and **Table ST4**). Interestingly, such relationship was particularly evident for the cingulate, inferior frontal gyri, and supplementary motor area.

## Discussion

In this study, we aimed to assess the brain prediction error indexed by the MMN elicited by cognitive and acoustic deviants inserted in a musical context. Results revealed that the neural sources of the MMNs were mainly localized in the auditory cortex. However, significant clusters of activity were also observed in the cingulate gyrus, hippocampal, and frontal areas. Notably, the contrast between cognitive versus acoustic deviants showed stronger activity within the cingulate gyrus for the cognitive deviants. Conversely, the acoustic deviants elicited stronger responses in the auditory cortex. At last, we revealed that musical expertise modulated the sources of the brain prediction error indexed by MMN to both categories of deviants. Notably, such modulation was stronger for cognitive deviants and involved especially the cingulate, inferior frontal gyri, and supplementary motor area.

The brain sources which generated the MMNs were coherent with the sources reported by previous literature. Specifically, several studies showed that auditory cortex, and especially Heschl’s gyrus together with superior and middle temporal gyri, is primarily implicated in the generation of the MMN ^9–11^. In addition, we detected sources in the medial and anterior cingulate gyrus, hippocampal areas, and frontal operculum/inferior frontal gyrus. This is also supported by previous research which highlighted frontal generators of MMN ^10^, proposing that they were necessary for the process of switching attention to the deviant stimulation ^15–18^.

Interestingly, we detected a dissociation between the sources underlying processing of cognitive versus acoustic deviants. Indeed, while auditory cortex was primarily recruited by the processing of acoustic deviants, a higher-order area such as the medial cingulate gyrus was stronger for the cognitive deviants. Moreover, insula and frontal brain areas such as anterior cingulate gyrus and inferior frontal gyrus exhibited greater activity than auditory cortex when observing the MMN sources of two of our cognitive deviants, namely rhythm modulation and transposition. In particular, the transposition deviant is thought to be the most cognitive deviant of the paradigm. Indeed, Putkinen and colleagues ^34^ showed that transposition was the only deviant not evoking larger MMN in music-trained children than in control ones. These findings broadened our understanding of MMN sources and auditory prediction error. In fact, there was no evidence in favour of a higher switching of attention for the cognitive versus acoustic deviants. Thus, the higher involvement of frontal brain regions observed in our study for the cognitive deviants should not be connected to the “attention switching” hypothesis ^15–18^ mentioned above. Conversely, we argue that cognitive and acoustic deviants elicit two diverse types of auditory prediction error. As a matter of fact, acoustic deviants are perceived as “mistakes” occurring in the music. On the contrary, cognitive deviants are actual changes of the musical information carried by the melodies. In other words, in the first case the brain may simply notice an impaired quality of the musical information, while in the second scenario, the prediction error operated by the brain would be more complex, leading to the understanding that musical information has actually changed. Notably, even though automatic and independent of a participant’s attention, the prediction error associated with changes in musical information were generated by higher-order brain areas usually associated with language processing and conscious cognitive abilities, such as inferior frontal gyrus ^62–64^ and cingulate gyrus ^65–67^. Conversely, our findings suggest that musical “mistakes” such as imprecise rhythms, small mistunings, or sudden variations in timbre would not require such complex processing. Indeed, in this case, the recruitment of the auditory cortex would be enough to detect the changes in the physical, acoustic features of the sounds.

Among our results, of particular interest is the role of cingulate gyrus which has been previously connected to several functions, including prediction error. For example, Alexander and colleagues ^68^ highlighted the role of anterior cingulate cortex (ACC) in processing behavioural error and signalling deviations between expected and observed events, describing it within the framework of reinforcement learning. Similarly, an activation likelihood estimation (ALE) meta-analysis investigated the neural correlates of prediction error in reinforcement learning. Authors found that ACC, medial prefrontal cortex (mPFC) and striatum were the key brain areas underlying prediction error, in studies that used both rewarding and aversive reinforcers ^69^. Another fMRI study investigated the brain activity underlying a numerical Stroop task, reporting activity in the ACC when participants processed errors in the task ^70^. Along this line, another contribution claimed that ACC learnt to predict error likelihood in each context, even for trials in which there was no error ^71^. A simulation study on mPFC and ACC provided modelling evidence in support of the role of these brain structures for error likelihood, signaling mistakes, and reward, concluding that they are central for learning and predicting the likely outcomes of actions whether good or bad ^72^. Furthermore, Bonetti and colleagues ^53,60,61^ showed that cingulate gyrus is of primary importance for both active encoding and recognition of auditory sequences, and that its involvement positively correlate with the strength of the recognition ^73^. Their findings revealed that the cingulate is more central within the whole brain network when encoding sounds than when resting ^53^. Moreover, they found that recognition of previously learned compared to novel melodies was associated with stronger cingulate activity ^60,61^. Along this line, a recent meta-analysis ^74^ on music perception, imagery and production highlighted the involvement of cingulate gyrus when participants were asked to do a variety of different tasks concerning music listening and production, and mental manipulation of sounds. Taken together, this evidence supports the idea that cingulate gyrus may be a key structure for extracting information from musical sequences and signalling variations from the previously learned melodies.

Conversely, acoustic “mistakes” involving basic acoustic features of musical sounds may recruit a more restricted network of auditory brain areas. This evidence is supported by previous studies employing simpler oddball and multi-feature paradigms which highlighted the primary role of auditory cortex in the MMN generations. For instance, Marco-Pallares ^11^ and colleagues reconstructed the main sources of MMN measured with EEG within supratemporal and middle temporal cortex, bilaterally. Similarly, Waberski and colleagues ^10^ found the main generators of MMN in supratemporal brain regions. They also reported secondary sources, with a longer latency, localized in the cingulum and right inferior temporal gyrus. Notably, this conclusion was reached even in intracranial electroencephalography (iEEG) recording, where MMN sources were observed in Brodmann areas 21 and 42, corresponding to middle temporal gyrus and posterior transverse temporal cortex, respectively ^75^. Moreover, additional evidence pointed out that the auditory cortex is mainly implicated in the processing of basic acoustic features of sounds and music. For instance, in a classic work, Zatorre and colleagues ^76^ argued that auditory cortices in the two hemispheres are specialized to extract fundamental acoustic features of both music and speech such as temporal and spectral content of sounds. Specifically, they reported that temporal resolution was better in left auditory cortical regions while spectral resolution of the sounds was greater in right auditory cortical regions. In a more recent review, King and colleagues ^77^ highlighted the complexity of the auditory cortex and its important role also for high-level cognitive processes. Still, they reiterated that auditory cortex shows selectivity for sound features, which is likely at the basis of processing of natural sounds, such as during speech and in real-life listening scenarios.

Finally, we revealed that musical expertise modulated the brain sources of the prediction error signal elicited by cognitive and acoustic deviants. Notably, this modulation was primarily evident in high-order brain areas such as the cingulate, inferior frontal gyri, and supplementary motor area. Moreover, this relationship was primarily evident for the cognitive deviants. This finding is coherent with a large corpus of studies which showed that the brains of musicians are different from non-musicians’. Indeed, the musician’s brain has been suggested as a model of neuroplasticity ^78^, being shaped by long-lasting musical training. This hypothesis was further supported by several longitudinal studies showing structural brain changes, especially in children, after exposure to musical training ^79,80^. Likewise, a recent meta-analysis revealed that structural and functional brain differences emerged when comparing the brain of musicians versus non-musicians ^81^. Back to MMN research, several works reported a stronger MMN activity recorded in brains of participants with higher musical expertise ^29,37–40^. Additionally, Vuust and colleagues ^29^ found different brain responses even across diverse categories of musicians. For instance, they revealed that the brain of jazz versus classical musicians was more sensitive to pitch and pitch-sliding deviants, features which are particularly involved in jazz training. In light of previous findings, our results provide additional evidence that musical expertise is associated with higher-level processing of music in the brain. Further, our study suggests that to outperform non-musicians when extracting varied information from musical melodies, musicians rely on stronger activity of higher-order brain areas such as cingulate and inferior frontal gyri, and supplementary motor area.

In conclusion, our study showed that the brain employs different strategies for processing cognitive and acoustic auditory prediction error, and that musical expertise modulates such mechanisms. Future research is called to investigate auditory prediction error in a wider array of cognitive and acoustic deviants and assess whether similar results arise when performing attentive tasks which require a conscious elaboration of the musical information. Moreover, as previously done by Tervaniemi and colleagues ^82^ and Pulvermüller & Shtyrov ^83^, future studies should investigate acoustic and cognitive deviants in contexts different from music, such as employing linguistic experimental design and investigating speech sound MMNs.

## Data availability

The codes are available at the following link: https://github.com/leonardob92/LBPD-1.0.git, while the multimodal neuroimaging data related to the experiment are available upon reasonable request.

## Acknowledgements

The Center for Music in the Brain (MIB) is funded by the Danish National Research Foundation (project number DNRF117). Moreover, we wish to acknowledge the financial support of the Academy of Finland (project number 133673).

LB is supported by Carlsberg Foundation (CF20-0239), Center for Music in the Brain, Linacre College of the University of Oxford, and Society for Education and Music Psychology (SEMPRE’s 50th Anniversary Awards Scheme).

Additionally, we thank the Italian section of *Mensa: The International High IQ Society* for the economic support provided to Francesco Carlomagno.

Finally, we thank the following persons for their help with MEG data collection: David Ellison, Anja Thiede, Chao Liu, Suvi Lehto, Jyrki Mäkelä, Simo Monto.

## Author contributions

Conceptualization: EB, MK, FC, LB, MT, MH, BG, SP; Methodology: LB, FC, EB; Software: LB, FC, NTH; Analysis: LB, FC, NTH; Investigation: MK, MT, MH, BG, SP, EB; Resources: EB, MT, MH, SP, PV, LB; Data curation: LB, FC, MK, EB; Writing - Original draft: LB, FC; Writing – Review & editing: LB, FC, EB, PV, MT, MH, MK, NTH; Visualization: FC, LB; Supervision: EB, PV, LB; Project administration: EB, PV, LB; Funding acquisition: PV, EB, MT, MH, LB, SP.

## SUPPLEMENTARY MATERIALS

As follows, supplementary materials related to this study. In the cases when the supplementary tables were too large to be conveniently reported in the current document, they have been reported in Excel files that can be found at the following link: https://drive.google.com/drive/folders/1GBrjTPTkf0JTbS1Q3PZ8_0zJyHfPm-bM?usp=sharing

### SUPPLEMENTARY TABLES

**Table ST1. MMN and MEG sensors**

Extended table showing the significant time-points over all MEG channels where the ERF elicited by the deviants was stronger than the one elicited by the standard sounds.

**Table ST2. Brain generators of MMNs**

Brain sources forming the significant cluster of activity reconstructed for the MMNs to the six deviants. For each voxel of the significant clusters, the table shows brain region label (from automated anatomical labelling (AAL) parcellation), hemisphere, t-value from the contrast deviant versus standard, and MNI coordinates.

**Table ST3. Brain generators of MMNs – Cognitive versus acoustic deviants**

Contrast between MMNs to cognitive versus acoustic deviants. For each voxel of the significant clusters, the table shows brain region label (from AAL parcellation), hemisphere, t-value from the contrast MMNs to cognitive versus acoustic deviants, and MNI coordinates.

**Table ST4. Cognitive and acoustic deviants and musical expertise**

Correlations between brain generators of MMNs to cognitive and acoustic deviants and participants’ musical expertise. For each voxel of the significant clusters, the table shows brain region label (from AAL parcellation), hemisphere, rho from the correlation between participants’ musical expertise and MMNs to cognitive or acoustic deviants, and MNI coordinates.

